# Multispecies phase diagram of biofilm architectures reveals biophysical principles of biofilm development

**DOI:** 10.1101/2021.08.06.455416

**Authors:** Hannah Jeckel, Francisco Díaz-Pascual, Dominic J. Skinner, Boya Song, Eva Jiménez-Siebert, Eric Jelli, Sanika Vaidya, Jörn Dunkel, Knut Drescher

## Abstract

Bacterial biofilms are among the most abundant multicellular structures on Earth and play essential roles in a wide range of ecological, medical, and industrial processes. However, general principles that govern the emergence of biofilm architecture across different species remain unknown. Here, we combine experiments, simulations and statistical analysis to identify shared biophysical mechanisms that determine biofilm architecture development at the single-cell level, for the species *Vibrio cholerae, Escherichia coli, Salmonella enterica*, and *Pseudomonas aeruginosa*. Our data-driven analysis reveals that despite the many molecular differences between these species, the biofilm architecture differences can be described by only two control parameters: cellular aspect ratio and cell density. Further experiments using single-species mutants for which the cell aspect ratio and the cell density are systematically varied, and mechanistic simulations, show that tuning these two control parameters reproduces biofilm architectures of different species. Altogether, our results show that early-stage biofilm architecture is determined by mechanical cell-cell interactions which are conserved across different species and, therefore, provide a unifying understanding of biofilm architecture development.

## Introduction

Bacterial biofilms are multicellular communities that grow on surfaces within a self-produced extracellular matrix [1, 2]. The composition and properties of this extracellular matrix varies widely between different species [3–5], but despite the molecular dissimilarities, biofilms of different species generally share a robustness against mechanical and chemical perturbations. Major research efforts over the past two decades [6–9] have established the ecological, biomedical and industrial importance of bacterial biofilms, and revealed that biofilms are highly abundant on Earth [10].

However, it is not well understood how multicellular properties of biofilms, such as mechanical stability, arise from the collective growth and spatiotemporal self-organization of these communities. Recent advances in live imaging techniques make it possible to observe the development of early-stage biofilms at single-cell resolution, starting from a single founder cell up to a few thousand cells [11–15].

Imaging-based studies have provided key insights into the importance of mechanical cell interactions [13, 16–23], cell surface attachment [18, 24–28], growth memory [15], external fluid flow [14, 29, 30], and the external mechanical environment [31–33] for the emergent architecture in biofilms. However, these studies were restricted to a single species and it remains an open question whether there exist common biophysical principles that govern biofilm architecture development across species.

To tackle this problem, we report here a combined experimental and theoretical investigation of three-dimensional (3D) biofilm architectures for the bacterial species, *V. cholerae, E. coli, S. enterica*, and *P. aeruginosa*. Each of these species displays different growth characteristics, extracellular matrix components, cell morphology, and biofilm architectures. To identify common architectural characteristics across different bacterial species, and to ultimately identify conserved biophysical principles for biofilm development, it is necessary to have quantitative metrics enabling comparisons between multicellular structures which are able to robustly distinguish different biofilm architectures. Building on recent tools for 3D biofilm image analysis [34], we are able to extract and quantify numerous single-cell properties and emergent collective properties from microscopy image data of individual biofilms. To analyze these measurements, we introduce here a statistical metric framework based on a general Chebyshev representation of the experimentally measured parameter distributions, which is able to distinguish different biofilm species based on their architectural features. This metric overcomes limitations of previous methods that relied on the assumption of normally-distributed data [13]. Since the underlying mathematical formulations of our analysis framework of 3D multicellular structures is generic, the method will be broadly applicable to other prokaryotic and eukaryotic multicellular structures in the future.

Through the quantitative biophysical analysis methodology outlined above, we find that emergent architectural differences across biofilms of different species correlate with variations in cell shape and local cell density. To test whether these correlations are due to causal relationships, we used mutants of a single species and particle-based computational modeling to independently explore the biophysical phase space of early-stage biofilm architectures. These experiments and simulations showed that two mechanical parameters (cell aspect ratio and the cell-cell attraction) jointly determine the emergent biofilm architecture across different species, which reveals a conserved principle for biofilm architecture development.

## Results and Discussion

### Quantifying early-stage biofilm architecture across species

To investigate the structural differences between biofilm architectures within and across bacterial species, we performed single-cell resolution imaging. For each of the four species *E. coli, V. cholerae, P. aeruginosa*, and *S. enterica*, 15 biofilms were grown in microfluidic flow chambers from a few surface-attached founder cells until they reached around 2000 cells, followed by imaging using confocal microscopy (Fig. 1A; Materials and Methods). Although all species formed colonies, the biofilm architectures of the four species were qualitatively different (Fig. 1A). To quantify the observed differences in biofilm shape and structure between species, we segmented all individual cells in all biofilms following Ref. [13]. Using the software tool BiofilmQ [34], we measured for each biofilm several single-cell properties such as cell length, cell diameter, and cell convexity, together with emergent collective properties, such as local cell number density and nematic order, resulting in a histogram for every one of the *m* = 16 measured properties (see S1 Text Section 1 for a complete list). Each biofilm is thus represented by a set of *m* histograms.

**Fig 1.**
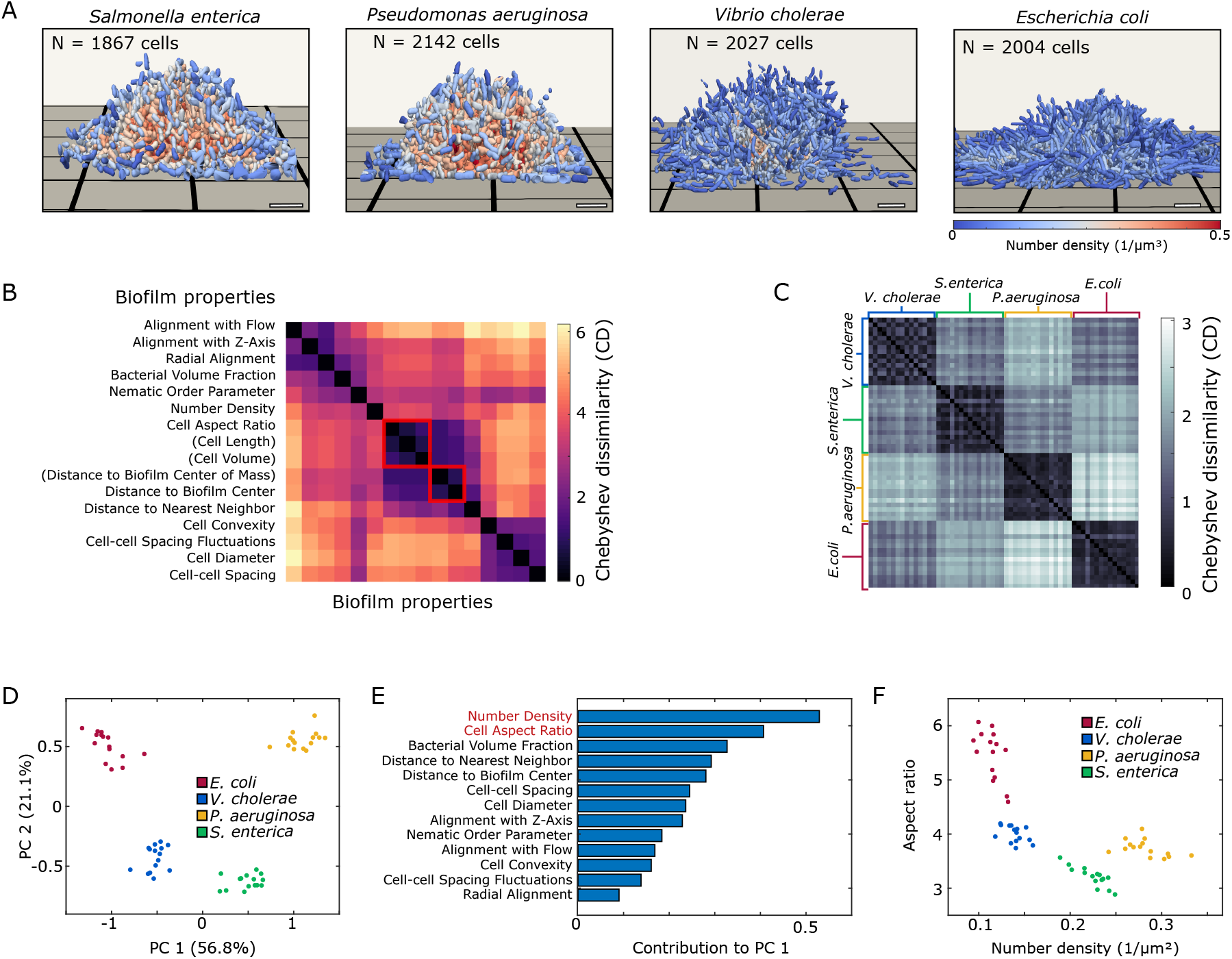
Early-stage biofilm architectures of different bacterial species can be quantitatively distinguished within a two-dimensional phase diagram derived from a statistical analysis of architectural properties. (A) Representative 3D biofilm architectures of four bacterial species reconstructed from segmented confocal microscopy images at comparable cell numbers (∼ 2000 cells). Each cell is colored according to the local density within its neighborhood, of radius 2 *μ*m. Scale bars, 5 *μ*m. (B) For each biofilm, we approximated the distributions of 16 measured properties with Chebyshev polynomials (S1 Text Section 1). Using the Chebyshev polynomials for each measured property, a Chebyshev dissimilarity (Cd) measure is defined (S1 Text Section 2). Using this measure, highly correlated properties are identified and reduced as indicated by red squares, leaving *p* = 13 relevant properties (S1 Text Section 3). (C) The Chebyshev dissimilarity also provides a robust and quantitative comparison of biofilm architectures from different species, as indicated by the block structure in this diagram. (D) Principal component analysis (PCA) based on the Chebyshev coefficient space (S1 Text Section 2) robustly distinguishes biofilms of the four different species *E. coli, V. cholerae, P. aeruginosa*, and *S. enterica*. (E) Cell aspect ratio and local number of neighbors are the key contributors to the first principal component (S1 Text Section 2). (F) Representing the experimental data in the mean aspect ratio vs. cell number density plane confirms that these two properties define a biophysically interpretable phase diagram to categorize biofilm architectures.

Previous approaches have used mean- and variance-based measures of these histograms [13] to distinguish biofilm architecture, however these measures do not carry information about the histogram’s shape and are therefore of limited utility. To broaden the scope of our statistical analysis and therefore the range of systems that it can be applied to, we sought a more general approach to systematically compare sets of histograms. To this end, we represented each empirically measured histogram with a Chebyshev polynomial of degree *d* = 20 using kernel density estimation (S1 Text Section 2). Replacing ∼ 2000 single cell measurements for each biofilm and each parameter with *d* + 1 = 21 polynomial coefficients allowed us to compress the experimentally observed data whilst retaining information about their distributions beyond mean values and variances. From a (*d* + 1) × *m* matrix containing all the Chebyshev coefficients for a given biofilm, we constructed a Chebyshev dissimilarity (Cd) measure, to compare two such matrices and hence two biofilms (S1 Text Section 2). Mathematically, Cd provides an upper bound on the cumulative *L*_1_-distance between collections of histograms.

Similarly, taking a vector of Chebyshev coefficients constructed from a single property across all biofilms, allows us to apply Cd to compare similarities of measured properties (S1 Text Section 2). To prevent double-counting, we identified highly correlated properties by performing clustering based on Cd and using the silhouette coefficient to determine the optimal cluster number (See S1 Text Section 2 for more details). This analysis left us with *p* = 13 essential properties which characterize biofilm architecture (Fig. 1B). When calculating Cd for each pair of biofilms using the 13 essential properties, we observe a robust distinction according to species, as evident from the block structure in Fig. 1C.

### Data-driven identification of the phase diagram of early-stage biofilm architecture

Principal component analysis (PCA) applied to the flattened (*d* + 1) × *p* = 21 × 13 dimensional vectors of Chebyshev coefficients representing each biofilm revealed that there are four distinct clusters corresponding to the four bacterial species (Fig. 1D). The information contained in the *p* = 13 distributions of measured parameters is therefore sufficient to capture the key architectural differences between species.

The first principal component, which explains more than 50% of the variation in the data, can be used as a scalar measure for biofilm architecture and will from here on be referred to as the biofilm architecture index (BAI). To investigate which of the measured properties could be responsible for the inter-species variation, we examined the contributions of each parameter to the BAI (Fig. 1E). The feature that contributed most to the BAI is the local cell number density, defined as the number of neighbors that a cell has within a 2 *μ*m radius. The second highest contributing feature was the cell aspect ratio. The prominent contributions of the cell number density and cell aspect ratio to the BAI suggest that variations in these two parameters across biofilms could be responsible for variation in the observed architectures. To verify that these two properties provide the basis for a suitable biophysical phase diagram of biofilm architecture, we plot each biofilm in the mean cell number density vs. mean cell aspect ratio plane (Fig. 1F). The clear separation of the four species in this two-dimensional phase space shows that biofilm architectures can be efficiently characterized by these two parameters. We note that classical liquid crystals can also be characterized by an aspect ratio vs. number density phase diagram [35, 36], which highlights an interesting analogy between passive nematic structures, and growth-active nematic biofilms.

### Altering biofilm architecture with cell aspect ratio mutants and cell-cell adhesion mutants

The four species analyzed in Fig. 1 differ in a large number of biological properties beyond the cell aspect ratio and number density. To test if cell aspect ratio and local density not only correlate with but also determine the different biofilm architectures observed across the four species, we generated several mutants in a single species, *V. cholerae*. By analyzing the biofilm architectures that arise from mutants within a single species, it is possible to isolate the effects of cell aspect ratio and local density on the biofilm architecture. To this end, we generated mutations in *mreB*, following Ref. [37], which resulted in different aspect ratios compared to the parental strain (Fig. 2A). For altering the cell aspect ratio, we preferred using *mreB* mutations instead of using antibiotics (such as cephalexin), because these mutations did not interfere with bacterial replication rates.

**Fig 2.**
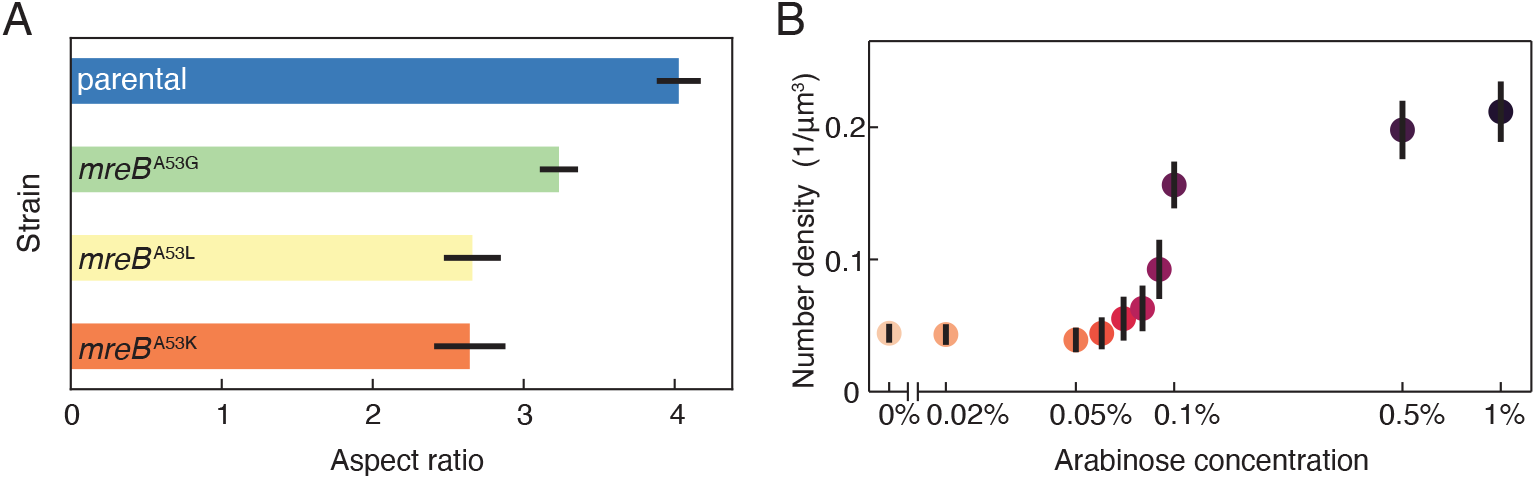
Cell aspect ratio and cell-cell adhesion can be precisely controlled in mutant strains. (A) Using different *V. cholerae* strains, the cell aspect ratio can be experimentally varied. Measurements were performed for *n* = 30 biofilms, including strains with point mutations in the *mreB* gene. Bar height corresponds to means, error bars indicate standard deviations. (B) Through additional mutations, the local number density can be varied experimentally, independent from the cell aspect ratio. Biofilms grown in the presence of different arabinose concentrations (*n* = 12 biofilms for each arabinose concentration), exhibit an increasing local number density with increasing arabinose concentration. Measurements were performed and averaged for Δ*rbmA* strains with either the wild type *mreB, mreB* ^A53K^, *mreB* ^A53L^, or *mreB* ^A53G^, harboring a plasmid with an arabinose-inducible *rbmA* expression construct (P_*BAD*_ -*rbmA*). Data points are colored according to the arabinose concentration, error bars indicate standard deviations.

To control the cell density, we introduced mutations that alter the abundance of the cell-cell attraction-mediating matrix protein RbmA [11, 13]; specifically, we deleted the native *rbmA* gene from the chromosome, and re-introduced a copy of *rbmA* under the control of a promoter that is inducible by the monosaccharide arabinose (Materials and Methods). By growing the cells in the presence of different levels of arabinose, we can therefore tune the level of the cell-cell attraction, resulting in different cell number densities (Fig. 2B). We introduced the *rbmA* mutation and inducible *rbmA* expression construct into the parental *V. cholerae* strain, as well as in strains with smaller aspect ratios, to perform a comprehensive experimental scan over the different cell aspect ratios and cell densities, which resulted in widely different biofilm architectures (Fig. 3A).

**Fig 3.**
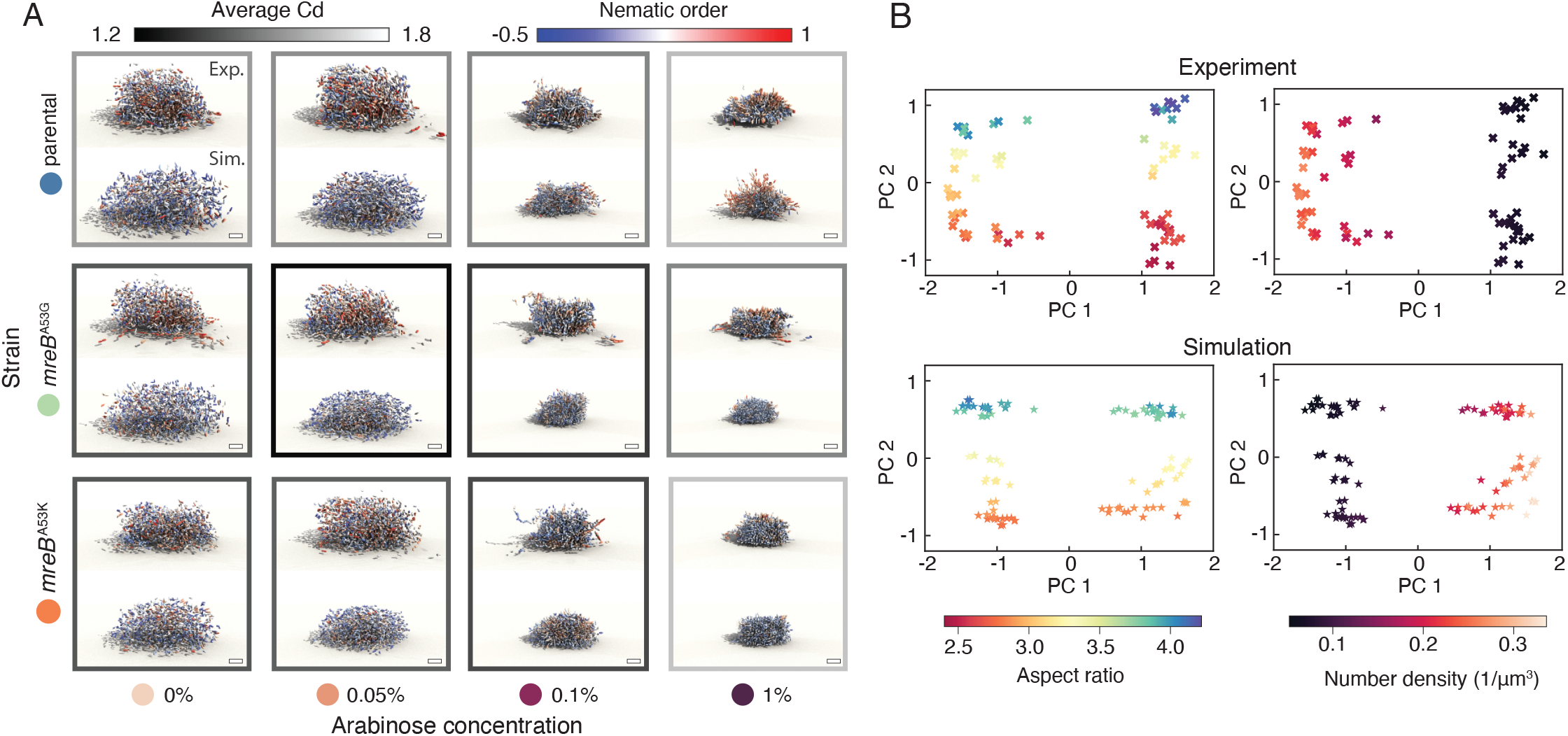
Experiments and simulations show that differences in biofilm architecture are driven by cell aspect ratio and cell-cell adhesion. (A) Renderings of *V. cholerae* biofilms (top, *N*_cell_ ∼ 2000) and corresponding best-fit simulations (bottom, *N*_cell_ = 2000) for combinations of three different mutants (arranged vertically) and four different arabinose concentration levels (arranged horizontally). Each cell in the biofilm is colored by the local nematic order around it. The outline of each grid panel is colored in grayscale by the average Chebyshev dissimilarity (Cd) between the corresponding experiments (*n* = 3) and the best-fit simulation (see Fig. S6 for the exact values). Scale bar, 5 *μ*m. (B) Two-dimensional PCA embedding of the Chebyshev features of *n* = 72 *V. cholerae* mutant biofilms (top), and a group of *n* = 114 simulations consisting of the top 5 best-fitting simulations (bottom) for each combination of strain and arabinose concentration. The PCA embedding is colored by average aspect ratio (left) and average local number density (right) of all the cells in each biofilm, confirming that these two parameters are principal determinants of biofilm architecture, consistent with Fig. 1E.

To understand whether the natural phase diagram of biofilm architectures for the different *V. cholerae* mutants is, like the phase diagram for the different species introduced in (Fig. 1F), also based on the cell aspect ratio and cell number density, we again performed PCA. Applying PCA to the vectors of Chebyshev coefficients for each biofilm and coloring the data points by aspect ratio (Fig. 3B, top left) and number density (Fig. 3B, top right) reveals that these parameters exactly correspond to the first two principal components of this embedding. Therefore, the appropriate phase diagram of biofilm architectures of *V. cholerae* mutants spanned by the aspect ratio and number density, consistent with the results for the different species in Fig. 1F.

### Computational model for biofilm growth based on mechanical interactions reproduces experimental biofilm architectures

Cell aspect ratio and cell-cell attraction, which were systematically varied for *V. cholerae* experimentally (Fig. 2A-B), are key parameters for the mechanical cell-cell interactions. To test if the effect of these parameters on the biofilm architecture is primarily due to changes in mechanical cell-cell interactions, we compared the experimental measurements for the *V. cholerae* strains with a computational model for biofilm growth in which cells only interact mechanically (Fig. 3A). In this model, which extends a previously introduced simulation framework [13, 14], individual cells are represented as growing, dividing ellipsoids which experience pairwise cell-cell interactions and cell-surface interactions that determine their overdamped positional and orientational dynamics. The cell-cell interactions account for both short-range steric repulsion together with RbmA mediated attraction [13, 14]. In addition to cell-surface steric repulsion [13, 14], our simulations now also include an effective cell-surface attraction to account for the surface attachment of *V. cholerae* before and during biofilm formation [38, 39]. To further refine the previously introduced minimal model [13, 14], we implemented strongly anisotropic friction effects to account for the fact that the matrix polymer network can suppress the transverse motions of cells [40–42] (S1 Text Section 3). We generally found that the inclusion of the cell anchoring to the bottom surface and the anisotropic matrix-mediated friction leads to a substantially improved agreement between experimentally observed and simulated biofilms (Fig. 3A), when comparing their architectural properties in terms of the Cd measure (S1 Text Section 3).

To compare the experimental biofilm architectures of the *V. cholerae* mutants with the computational model, we performed systematic parameter scans to identify the values of simulation parameters which correspond to a given experimental system. Specifically, we performed > 6, 000 simulations to search the parameter space of cell length at the time of division, range of cell-cell repulsion force, range of cell-cell attraction force, and strength of the cell-cell attraction (S1 Text Section 3), with the remaining parameters determined from a previous experimental biofilm calibration [13, 14] (see S1 Text Table S2). The best-fitting parameter values for a given experiment were determined by taking the values with the smallest Cd between experiment and simulation (Fig. S5). Using the fitted parameter values, we see a qualitative agreement between the experiment and simulation across various combinations of cell aspect ratio mutants and arabinose concentration levels (Fig. 3A). This agreement between the biofilm architectures obtained from the experimental parameter scan and the simulation parameter scan indicates that changes in cell aspect ratio and cell-cell attraction cause changes in the biofilm architecture through their effects on mechanical cell-cell interactions.

Analogous to our analysis of experimental biofilm data from *V. cholerae* mutants (Fig. 3B, top row), we again applied PCA to the Chebyshev coefficients of *n* = 114 simulated biofilms and colored the points according to aspect ratio (Fig. 3B, bottom left) and number density (Fig. 3B, bottom right). Consistent with the experimental results, these diagrams reveal that the principal component axes correspond to the number density and aspect ratio, respectively. Similar to the results for the different species (Fig. 1F) and the *V. cholerae* mutants (Fig. 3B, top row), the PCA for the simulations (Fig. 3B, bottom row) indicates that the appropriate phase diagram of biofilm architectures is spanned by the aspect ratio and number density.

### Biofilm architecture of one species can be transformed into architecture of another species by changing mechanical control parameters

Given that the cell aspect ratio and number density in biofilms are the key control parameters for the biofilm architecture, we sought to understand which emergent architectural features change in the aspect ratio – density phase plane, and which conclusions can be drawn from these changes. We therefore plot the experimental biofilms for the different species and *V. cholerae* mutants together with our simulation results in the aspect ratio – density phase plane (Fig. 4), and color-code different emergent properties of the biofilm architecture in each panel: Fig. 4A shows the BAI, and panels B and C show the nematic order fluctuations and the biofilm surface area per volume, respectively. The nematic order fluctuations and the biofilm surface area per volume are both independent from our statistical analysis, which ensures that our observations are not a particularity of the BAI but also reflected in other biofilm-architecture related measures. The graphs in Fig. 4 show that number density is the key contributor to biofilm architecture, and cell aspect ratio has a more subtle influence.

**Fig 4.**
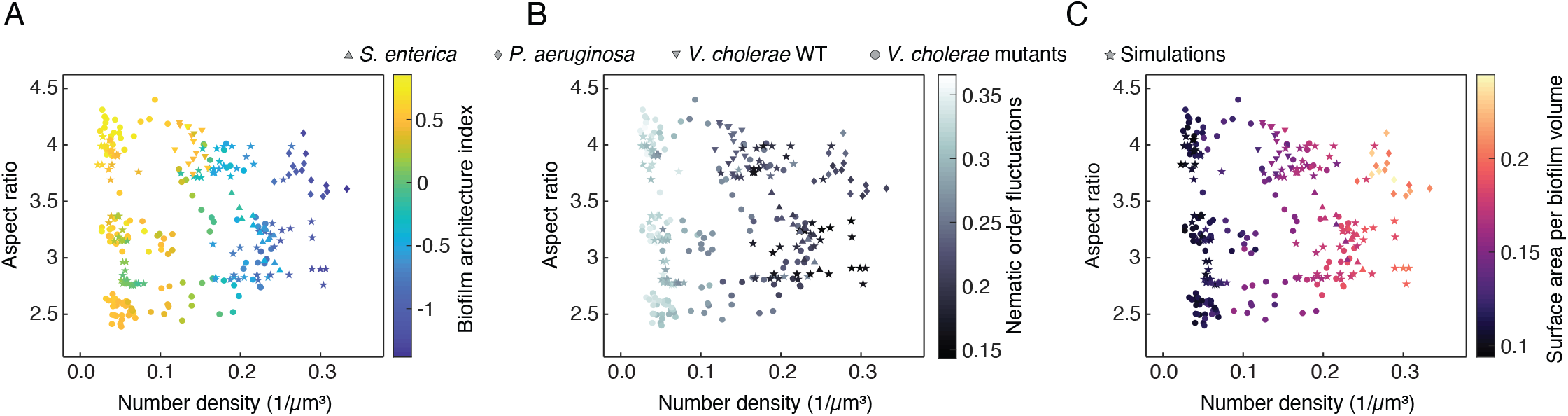
Joint phase diagram combining experimental biofilms from different species shown in Fig. 1A with the experimental biofilms of *V. cholerae* mutants, and the and simulated biofilms from Fig. 3. Each biofilm in the phase diagram is (A) colored by the biofilm architecture index BAI, (B) colored by the an emergent collective property of the biofilm architecture, the variance of the nematic order parameter, (C) colored by another emergent collective property, the surface area per biofilm volume.

The emergent properties of the biofilm architecture in Fig. 4 for the experimental and simulated biofilms agree very well for all regions in the phase diagram, indicating that the mechanics-based simulations capture the emerging biofilm architecture, irrespective of the particular species under investigation. Even though the specific molecular structure and composition of the extracellular matrix differ widely for the different species, these molecular details only indirectly influence this phase diagram through the number density.

Finally, the phase diagrams in Fig. 4 show that while the data points from each species inhabit a particular region in the phase plane, the *V. cholerae* mutants spread across the phase plane regions of different species. In each phase plane region, the emergent properties of the biofilm architecture of the *V. cholerae* mutants match those of the particular species inhabiting this phase plane region. These results show that the biofilm architecture of *V. cholerae* can be modified to reproduce the biofilm architecture of other species, by simply tuning the control parameters of the phase diagram (cell aspect ratio and cell number density).

## Conclusions

By performing single-cell resolution imaging on early-stage bacterial biofilms of several bacterial species, we found that the emergent biofilm architecture correlates with differences in cell aspect ratio and local cell number density. By systematically varying the aspect ratio and cell-cell attraction using mutants of a single bacterial species, we then showed that these parameters determine the observed architectural differences.

Extensive particle-based simulations of biofilm growth support this conclusion and further revealed that the impact of these parameters on the emergent biofilm architecture reflects the underlying effective mechanical cell-cell interactions. Our combined experimental and theoretical results show that bacterial biofilm architectures populate an aspect ratio – number density phase diagram, similar to classical liquid crystals. By changing the cell aspect ratio and number density of a particular species, this species can reproduce biofilm architecures of other species, even though the extracellular matrix composition and cellular properties can differ widely between species.

## Materials and methods

### Bacterial strains and media

All *V. cholerae* strains used in this study are derivatives of a rugose variant of the O1 biovar El Tor wild type strain N16961 [43]. The *E. coli* strain used in this study (KDE2011) is a derivative of the AR3110 wild type [44], carrying a point mutation in the promoter of the gene *csgD*, which increases biofilm formation [45]. The *S. enterica* strain used here (KDS38) is a derivative of the UMR1 wild type [46], carrying a mutation in the promoter of *csgD* (formerly called *agfD* in *Salmonella*), which increases biofilm formation [47]. The point mutations in the *E. coli* and *S. enterica* strains were necessary to grow isolated biofilm colonies in our experimental conditions. The *P. aeruginosa* strain used here (KDP63) is a derivative of the PAO1 wild type [48] (obtained from Urs Jenal, Basel). The *V. cholerae, E. coli*, and *S. enterica* strains carried a plasmid driving the production of sfGFP using the P_*tac*_ promoter. The *P. aeruginosa* strain KDP63 carried a high-copy number plasmid producing the fluorescent protein YPet under the control of a pX2 promoter [49].

To engineer *V. cholerae* strains with a different cell length and width, amino acid 53 of the native MreB protein was replaced according to Monds *et al*. [37]. These modifications were introduced to the chromosome of *V. cholerae* by conjugation using the *E. coli* strain S17-1 *λpir* [50] and the pKAS32 suicide vector [51], containing *mreB* with the corresponding mutation and 500 bp upstream and 500 bp downstream from the codon that codes for amino acid 53 of MreB. To control the expression of *rbmA* in *V. cholerae*, inducible strains were created by conjugating a plasmid that contained P_*tac*_*-sfGFP* and P_*BAD*_*-rbmA* constructs. This plasmid enabled us to vary the production of RbmA by adding different concentrations of arabinose to the growth medium [13]. All strains, plasmids, and oligonucleotides that were used in this study are listed in Table S1, Table S2, and Table S3, respectively.

For overnight cultures or strain construction, cells were either grown in liquid Luria–Bertani–Miller broth (LB-Miller; 10 g L^−1^ tryptone, 5 g L^−1^ yeast extract, and 10 g L^−1^ NaCl) or LB-Miller without salt (10 g L^−1^ tryptone and 5 g L^−1^ yeast extract) with the corresponding antibiotic and shaking at 250 rpm, or using agar-solidified LB media (containing 1.5% agar). All *V. cholerae* biofilm experiments were performed in M9 minimal medium, with the following composition: M9 minimal salts (M6030, Sigma), 2 mM MgSO_4_, 100 μM CaCl_2_, MEM vitamins, 0.5% glucose, 15 mM triethanolamine (pH 7.1), and gentamicin (30 μg mL^−1^). *E. coli* biofilm experiments were performed in tryptone broth (10 g L^−1^ tryptone) supplemented with kanamycin (50 μg mL^−1^). *S. enterica* biofilm experiments were performed in tryptone broth supplemented with spectinomycin (100 μg mL^−1^). *P. aeruginosa* biofilm experiments were performed in FAB medium, with the following composition: CaCl_2_ (11 mg L^−1^), MgCl_2_ (93 mg L^−1^), (NH_4_)_2_SO_4_ (2 g L^−1^), Na_2_HPO_4_·2H_2_O (6 g L^−1^), KH_2_PO_4_ (3 g L^−1^), NaCl (3 g L^−1^), glucose (25 ml L^−1^), and the trace metals solution (100 ml L^−1^). The trace metals solution consists of CaSO_4_·2H_2_O (2 mg L^−1^), FeSO_4_·7H_2_O (2 mg L^−1^), MnSO_4_·H_2_O (0.2 mg L^−1^), CuSO_4_·5H_2_O (0.2 mg L^−1^), ZnSO_4_·7H_2_O (0.2 mg L^−1^), CoSO_4_·7H_2_O (0.1 mg L^−1^), NaMoO_4_·H_2_O (0.1 mg L^−1^), and H_3_BO_3_ (0.05 mg L^−1^).

### Flow chamber biofilm experiments

Biofilms were grown in microfluidic flow chambers, which were made from polydimethylsiloxane bonded to glass coverslips using an oxygen plasma, with four to eight identical flow channels on a single coverslip. All flow rates were controlled using a syringe pump (PicoPlus, Harvard Apparatus). The microfluidic channels were 500 μm wide and 7 mm long. For *V. cholerae, E. coli*, and *S. enterica*, channels with height 100 μm were used, whereas for *P. aeruginosa*, channels with height 300 μm were used. Each biofilm is considered as a biological replicate.

For *V. cholerae* biofilm growth, overnight cultures grown in liquid LB-Miller with gentamicin (30 μg mL^−1^) at 28 °C were diluted 1:200 into fresh LB-Miller with gentamicin and grown for 2 h. Then, these cultures were adjusted to an optical density at 600 nm (OD_600_) of 0.001 and used to inoculate a microfluidic channel. The cells were given 1 h at room temperature to attach to the glass surface without flow, before fresh M9 medium with gentamicin was flown through the channel at a rate of 50 μL min^−1^ for 45 s, to wash away the non-attached cells. Then, the flow rate was set to 0.5 μL min^−1^ for the remainder of the experiment, and the flow channel as incubated at 25 °C.

For *E. coli* biofilm growth, overnight cultures were grown in liquid LB-Miller with kanamycin (50 μg mL^−1^) at 37 °C. These cultures were diluted 1:2000 into tryptone broth and used to inoculate a microfluidic flow chamber. The cells were given 1 h to attach to the substrate without flow, before washing away non-adherent cells using tryptone broth with kanamycin at a flow rate of 50 μL min^−1^ for 45 s. Then, the flow rate was set to 0.1 μL min^−1^ for the remainder of the experiment, and the flow channel was incubated at 25 °C.

For *S. enterica* biofilm growth, overnight cultures were grown at 37 °C in liquid LB-Miller without salt, supplemented with spectinomycin (100 μg mL^−1^). The overnight cultures were diluted 1:2000 and used to inoculate a flow channel. After giving the cells 1 h to attach to the coverslip without flow, the non-attached cells were washed away with tryptone broth supplemented with spectinomycin for 45 s using a flow rate of 50 μL min^−1^. The flow rate was then set to 0.1 μL min^−1^ for the remainder of the experiment, and the flow channel was incubated at 25 °C.

*P. aeruginosa* strains were grown overnight in 5 ml liquid LB-Miller with 30 μg mL^−1^ gentamicin at 37 °C with shaking. The overnight culture was back-diluted 1:200 in 3 mL LB-Miller and grown until OD_600_ = 0.5. This culture was subsequently diluted 1:1000 in FAB medium and used to inoculate microfluidic flow chambers. After allowing cells to attach to the glass coverslip for 1 h at 30 °C without flow, the cells were washed for 50 s using a flow rate of 200 μL min^−1^. The flow rate was then set to 3 μL min^−1^ for the remainder of the experiment, and the flow channel was incubated at 30 °C.

### Image acquisition

Biofilms were imaged using a electron-multiplying charge-coupled device camera (EMCCD, iXon, Andor) and a Yokogawa confocal spinning disk unit mounted on a Nikon Ti-E inverted microscope, and an Olympus 100 × silicone oil (refractive index = 1.406) objective with a 1.35 numerical aperture. The fluorescent protein sfGFP was excited using a 488 nm laser. Three-dimensional images were acquired during biofilm growth every 60 min, using a *z*-spacing of 400 nm. The hardware was controlled using Matlab (Mathworks). A live feedback between image acquisition, image analysis, and microscope control was used to automatically detect the biofilm and expand the imaging field during growth in 3D, as described by Hartmann *et al*. [13], to minimize the laser exposure of the growing biofilm.

## Supporting information

**S1 Text. File containing supporting information and supplementary figures**. (PDF)

**S1 Table. Bacterial strains used in this study**. (PDF)

**S2 Table. Plasmids used in this study**. (PDF)

**S3 Table. DNA oligonucleotides used in this study**. (PDF)

## Acknowledgements

We are grateful to Lucia Vidakovic for generating several strains that were used in this study, and to Urs Jenal and Benoit-Joseph Laventie for *P. aeruginosa* strains. This research was supported by grants from the Studienstiftung des deutschen Volkes and Joachim Herz Stiftung (to H.J.), a Mathworks Fellowship (to D.J.S.), the MIT Mathematics Robert E. Collins Distinguished Scholar Fund (J.D.), the European Research Council (StG-716734), Deutsche Forschungsgemeinschaft (DR 982/5-1), Minna James Heineman Foundation, Bundesministerium für Bildung und Forschung (TARGET-Biofilms), and the National Center of Competence in Research AntiResist funded by the Swiss National Science Foundation (grant number 51NF40 180541).

## Author contributions

H.J., F.D-P., D.J.S., and B.S. contributed equally to this work. K.D. and J.D. designed and supervised the project. F.D-P. performed experiments with all species except *P. aeruginosa*. E.J.S. performed some of the experiments with *V. cholerae*. E.J. and S.V. performed experiments with *P. aeruginosa*. B.S. performed the particle simulations. H.J., F.D-P., D.J.S. and B.S. performed data analysis. All authors wrote the paper.

